# Structural view on the role of WT1’s zinc finger 1 in DNA binding

**DOI:** 10.1101/284489

**Authors:** Raymond K. Yengo, Elmar Nurmemmedov, Marjolein M Thunnissen

## Abstract

The WT1 protein is a transcription factor that controls genes involved in cell proliferation, differentiation and apoptosis. It has become increasing apparent that WT1 can act both as a tumor suppressor and oncogene in a tissue specific manner. This opposing role of WT1 is linked to its underlying transcriptional regulatory function, which involves the specific binding to its regulatory elements on gene promoters. WT1 binds DNA using it C-terminal domain made up of 4 C2H2-typ zinc fingers. This same zinc finger domain is used to bind RNA and proteins and it is still not clear how each zinc finger contributes to this promiscuous binding behavior. The molecular details of DNA binding by zinc finger 2 to 4 have been described but it remains to be determined whether or not zinc finger 1 binds DNA and if so whether it exhibits any DNA binding specificity. We present the X-ray structures of zinc finger 1 to 3 bound to a 9 bp and an 8 bp DNA. The two structures refined to 1.7 Å, show no DNA binding specificity for zinc finger 1. The only DNA interactions involving zinc finger 1 are crystal-packing interactions with a symmetry related molecule. In the structure of zinc finger 1 to 3 bound to the 9 bp DNA we observe a shift in the DNA binding positions for zinc fingers 2 and 3. These structures provide molecular detail into the WT1-DNA interaction showing that zinc finger 1 only modestly contributes to DNA binding affinity through transient interactions. The dislocation of zinc finger 2 and 3 emphasizes the importance of zinc finger 4 for maintaining gene transcriptional specificity.

## Introduction

The WT1 protein, a product of the Wilms’ tumour 1 gene (*WT1*) is a zinc finger transcription factor implicated in a number of cellular processes particularly the development of the urogenital system and nervous system [1, 2]. This gene was originally cloned in the early 1990s as a candidate factor for the childhood kidney malignancy commonly known as Wilms’ tumour. Wilms tumour occurs with a frequency of about 1 in 10,000 mostly affecting children usually under the age of 8 [3]. Genetic analysis of Wilms’ tumour patients have found destabilizing mutations in 10% of sporadic Wilms’ tumour cases resulting in the conclusion that WT1 is a tumour suppressor [4, 5]. Mutations in this gene are also responsible for other diseases such as the Denys-Drash, Frasier and WAGR syndrome [6-8]. Interestingly the *WT1* gene has also been described as an oncogene in certain cancers such as leukemia because of the fact that elevated expression of wild-type WT1 is observed in these tumours [9-11].

There exist four major isoforms of WT1 as a result of 2 splicing events. The first includes a 17 amino acid peptide in exon 5 and the second include the tripeptide, lysine, threonine and serine (KTS) at the 3’end of exon 9. Many other minor isoforms of WT1 have been described, resulting in a large repertoire of WT1 isoforms presently numbering about 36 [12-16]. The WT1 protein consists of an N-terminal regulatory domain and a C-terminal recognition and binding domain. The N-terminal domain is mainly a non-structured proline-glutamine rich domain containing the activation and repression regulatory regions essential for protein-protein interactions [17]. The C-terminal domain is composed of 4 Krüpple-like C2H2 zinc fingers separated by canonical linker sequences, TG(E/V)KP that is mostly conserved except in the case of the +KTS isoforms where the linker region between zinc finger 3 and 4 is longer. This C-terminal domain is primarily a DNA binding domain but it also binds RNA and proteins. Post-translational modifications have resulted in some modified forms of WT1. There is the sumoylation of two lysines (K73 and K177) at the N-terminal part of the protein [18]. The particular significance of this post-translational sumoylation is not yet known. There is also another modification at the C-terminal part of the protein, which is phosphorylation of two serine residues (S365 and S393) in zinc fingers 2 and 3. Phosphorylation of DNA binding domains is a regulatory mechanism used to modulate the activity of transcription factors [19].

WT1 is a transcription factor with over 20 known targets, acting as an activator or repressor of gene transcription. The molecular mechanism that selects the targets and decides the type of regulation is not yet characterized, and it is believed to be isoform as well as cell specific. In order to understand this mechanism, the DNA binding activity, exclusive to the C-terminal, zinc finger domain has been extensively studied [20, 21]. The WT1 high affinity binding site, WTE with the nucleotide sequence, 5’GCGTGGGAGT3’ has been identified [22]. This and other studies, using methods such as oligonucleotide libraries screening, DNase footprinting, whole genomic PCR and bacteria on hybrid screens has led to the consensus WT1 DNA binding site used today. Subsequent studies have identified a stronger binding to a series of longer sites summarized as 5’GCG(T/G)GGG(C/A)G(T/G)(T/A/G)(T/G)3’ [22-26]. This sequence and the WTE sequence are very similar to the early growth receptor 1 (EGR-1) consensus sequence, 5’GCGGGGGCG3’. WT1 zinc fingers 2 to 4 share a 65% sequence identity with zinc fingers 1-3 of EGR-1. A recent structure of the C-terminal zinc finger domain of WT1-KTS isoform in complex with DNA has been reported [27]. This structure in combination with an NMR structure shows the specific interaction of the zinc fingers 2-4 with DNA. The three C-terminal fingers are wrapped around the DNA in a manner reminiscent of the interaction between the Zif268 zinc fingers with its cognate DNA [28]. Zinc finger 1 however does not bind in the major groove and makes no specific interactions with the DNA. Zinc finger 1 however makes contact with the DNA backbone.

Sequence alignment (**Figure 1**) of the four ZFs of WT1 reveals that ZF1 differs substantially from the other three zinc fingers. It has been observed that the amino acids at positions –1, 2, 3, and 6 of the α-helix within a zinc finger are responsible for determining the DNA-binding specificity [29]. However the amino acids found at the recognition positions in ZF1 are unconventional for DNA binding C2H2 zinc fingers. This DNA-binding amino acid sequence discrepancy for ZF1, the crystal structure which shows that ZF1 does not bind in the major groove of the target DNA and the inability of several studies to identify a consensus binding sequence for ZF1 suggest that zinc finger 1 may not be a DNA binding zinc finger or may only bind DNA nonspecifically. In fact, an alternative role for zinc finger 1 in RNA binding has been proposed [30]. This however does not eliminate the possibility of DNA binding by ZF1. Some authors find evidence for a reduction in the DNA binding activity of ZF1-deleted WT1 constructs by as much as 90% [22] while orders observe less dramatic effects [25, 26, 31, 32]. As a result of the inconclusive evidence as to whether ZF1 is a DNA binding finger or not, present sequence analyses to identify WT1 target genes use the minimal WTE sequence. This could be misleading if ZF1 is indeed a specific DNA binding zinc finger. For a complete understanding of the DNA binding properties of WT1, more evidence is needed to establish the DNA binding role of WT1 ZF1. Here we report the crystal structures of zinc fingers 1-3 of WT1 bound to a 9 bp and to an 8 bp DNA binding site. The structures show conclusively that zinc finger 1 does not bind DNA or at the most only binds DNA transiently and that zinc finger 2 and 3 bind DNA with some plasticity as they are unexpectedly shifted in these structures. This data will enable a clearer understanding of the transcriptional role of WT1.

**Figure 1:**
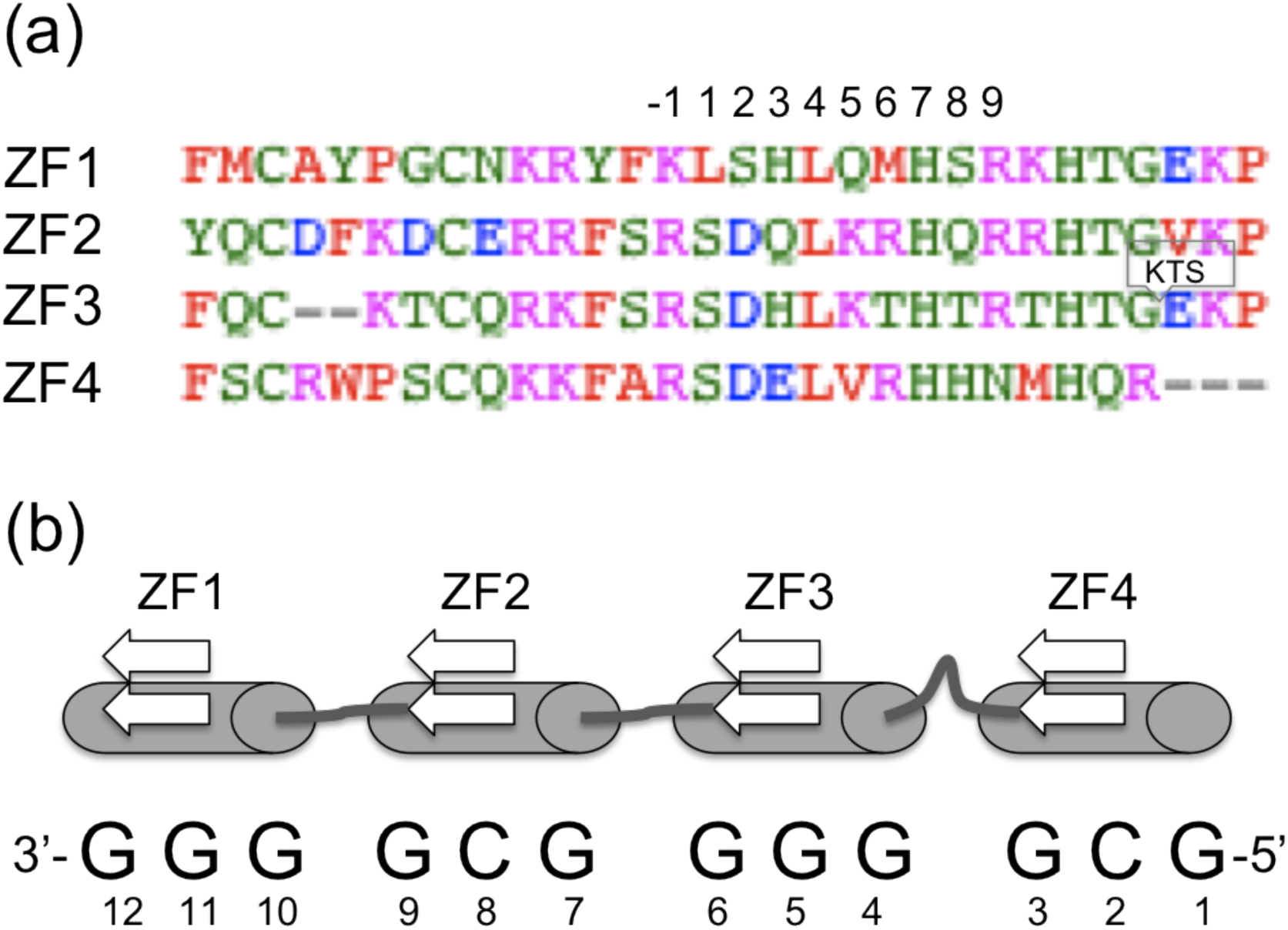
(a) Sequence alignment of WT1 zinc fingers 1 to 4. The amino acid positions in the helix are numbered according to the convention for DNA binding C2H2 zinc fingers. Differences between zinc finger 1 and the rest are evident at the DNAbindingpositions--1,2and6.(b)SchematicrepresentationoftheWT1zinc finger domain showing its DNA binding site and the numbering of the coding strand used in this study.

## Materials and Methods

### Sample production, activity assay and complex preparation

Six truncations of the WT1 C-terminal domain were used in this study, including ZF14-, ZF14+ and ZF24-, ZF24+, ZF13 and ZF23. The numbers 23, 13, 24 and 14 are used to represent zinc fingers 2 to 3, 1 to 3, 2 to 4 and 1 to 4 respectively. The + and – signs indicate the presence or absence of the KTS insert. Sample handling in the step before crystallizations was as previously described [33]. The cloning of all the 6 truncations used in this study was the same, but for the use of a primer exclusive for the 3’ end of the ZF23 and ZF13 cDNA. The primers sequences used for the ZF13 and ZF23 were as follows;

ZF13 Forward: 5 ’ ccatggagaaacgccccttcatgtgtgctta 3’

Reverse: 5’-ctaacctgtatgagtcctggtgtgggt-3’

ZF23

Forward: 5’ ccatggagaaaccataccagtgtgacttc 3’

Reverse: 5’-ctaacctgtatgagtcctggtgtgggt-3’

The expression and purification of the ZF13 and ZF24 was exactly as described for the ZF24 and ZF14 constructs. The cells were grown at 37°C, induced with 0.5 M IPTG, harvested, lysed and the protein was purified by a combination of ion exchange and gel filtration chromatography. Purified samples were analysed by SDS gel electrophoresis and dynamic light scattering. The Electrophoretic Mobility Shift Assays (EMSA) was performed to ascertain the DNA binding activity of the different WT1 zinc finger truncations.

The 2 double stranded DNA sequences used in this study are shown in **Figure 1**. The 2 pairs of single stranded oligonucleotides were chemically synthesized by Tag Copenhagen and shipped as lyophilized reverse phase cartridge purified DNA. Each oligonucleotide was re-suspended in 10 mM Tris, pH 7.0 to a final concentration of 200 µM. The complementary oligonucleotides were mix in a 1:1 molar ratio, heated to 95°C for 15 min and slowly cooled to room temperature to obtain 100 µM of double stranded DNA. The DNA duplex was loaded onto a HiTrap Q FF column (GE Healthcare) pre-equilibrated in 10 mM Tris buffer (pH 7.0), and eluted with a linear NaCl gradient to separate the duplex from any extra single stranded DNA in the annealing mixture.

Two complexes were prepared in this study including the complex between the WT1 ZF13 and a 9 bp DNA (ZF13/9bp) and the complex between the same ZF13 and an 8 bp DNA (ZF13/8bp). The sequence of the DNA used is presented in **Figure 2** and represented the binding site for zinc fingers 1 to 3 extracted from the complete WT1 DNA binding site. To obtain the complex, double stranded DNA was gradually titrated into the protein mixture to a final protein:DNA stiochiometry of 1:1.2 in binding buffer containing 20 mM Tris-HCl pH 7.0, 150 mM KCl, 1mM MgCl_2_, 10 µM ZnCl_2_, 1 mM DTT and 1 mM PMSF making sure not to exceed a complex concentration of 5 μM. KCl was further added to a final concentration of 300 mM to keep the complex in solution. The complex was then concentrated and purified on a Superdex 75 gel filtration column in the same binding buffer containing 300 mM KCl. The fractions containing the complex were filter concentrated, quantified and used for crystallization.

**Figure 2:**
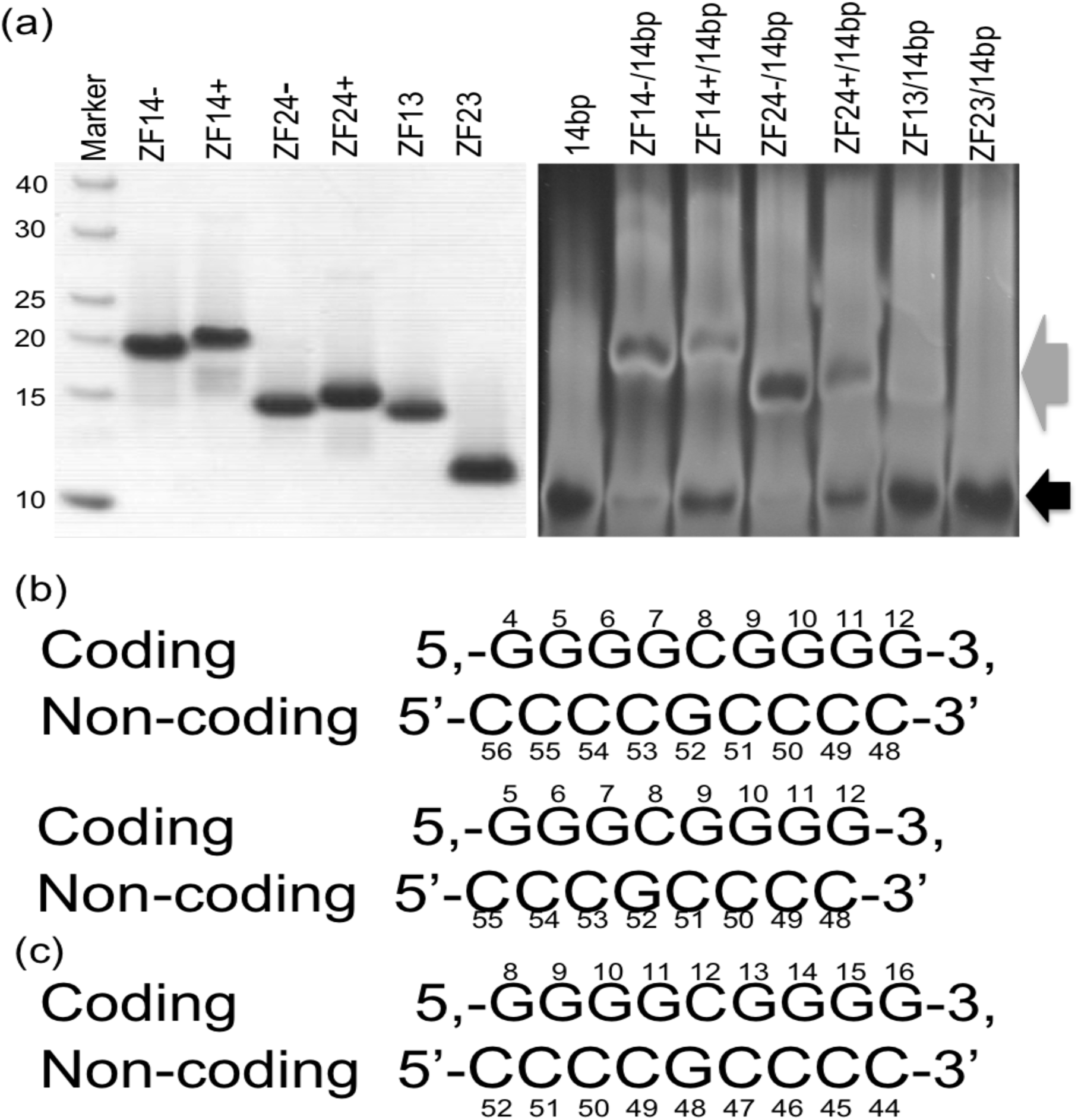
(a) Chromatograms showing the migration of the protein and complexes on gel. The first panel shows an SDS gel of the purified zinc finger truncations. The second panel shows the migration of the complex and free probe I an EMSA experiment. The grey arrow indicates position of complex and the black arrow, the position of free probe. (b) The 9 bp and 8 bp DNA sequences used in the crystallization. (c) The renumbering of the 9 bp to reflect its binding position in the crystal structure. This is the numbering used to describe the structure.

### Crystallisation, structure determination and analyses

The same protocol was used for the complex formation, crystallization, data collection and analysis of the two complexes presented in this study. Crystals were grown using the hanging drop vapor diffusion method at 15°C. About 5 hits were obtained from the initial sparse matrix screen, the crystallization Kit for DNA (Sigma Aldrich) but only the most promising were optimized. The drop contained 1 µL of complex and 1 µL of reservoir solution consisting in one case of 50 mM cacodylate pH 6.0, 10 mM MgCl_2_, 2 mM spermine, 10 mM CaCl_2_ and 12% isopropanol and in the other of 50 mM cacodylate pH 6.5, 15 mM MgCl_2_, 2 mM spermine, 20 mM CaCl_2_ and 11% 2-propanol. Crystals appeared after 1-2 days and grew to their full size of about 100 × 100 × 250 μm^3^ in one week. The crystals were transferred to a cryo solution containing the reservoir solution and 20% glycerol and flash cooled in liquid nitrogen. Data was collected at the Cassiopeia I911-3 beamline at the MAX-lab synchrotron radiation facility in Lund, Sweden. The diffraction data was integrated, scaled and converted to structure factors using the XDS data processing package [34]. The ZF13/9bp structure was solved by MRSAD (Molecular Replacement combined with Single anomalous Scattering) using the program Phaser-EP from the Phenix program suite [35] and initial phases from both the WT1 ZF14-14bp structure (PDB code 2PRT) and the WT1 ZF24-9bp structure [27]. The ZF13/8bp structure was solved by molecular replacement [36] with Phaser from the CCP4 program suite using the same search models as for the ZF13/9bp [37]. Initial model building into the electron density map was done manually in Coot [38]. Rigid body refinement was performed followed by multiple steps of restrained refinement using TLS with Refmac from the CCP4 suite [39]). The TLS groups were chosen such that each zinc finger, DNA strand and the zinc atoms each formed a unique group. Evaluation of model fit to density, model building and correction as well as the addition of waters was performed using Coot [38]. The final structure refinements were carried out using Phenix-refine [40]. Molecular graphic figures were prepared with PyMOL [41].

## Results

### Sample preparation

All the protein samples used in this study had a purity and homogeneity of more than 90% verified by gel electrophoresis and dynamic light scattering. Comparative DNA binding activity, using an electrophoretic mobility shift assay were the gels were stained with the DNA stain, SYBR Green (Invitrogen), indicated that ZF13 binds DNA less effectively than its ZF14 and ZF24 counterparts but more effectively than ZF23 (**Figure 2**) These results are consistent with earlier results supporting a DNA binding role for zinc finger 1 [25].

### Structure of WT1 ZF13 bound to its 9 bp cognate DNA (ZF13/9bp)

Crystals were grown of ZF13, a truncation of the WT1 zinc finger domain including zinc fingers 1-3 (residues MET 319-407) in complex with its 9 bp cognate DNA. Data was collected on the crystals of this complex (ZF13/9bp) and indexed using the XDS program package in space group P4(3)2(1)2 (**Table 1**) [34]. The phases were obtained by a combination of single anomalous dispersion and molecular replacement [35] with 1 molecule per asymmetric unit and a solvent content of 40.1%. The structure of ZF1 and ZF23 from the PDB entry, 2PRT and the structure of the 9 bp DNA built from sequence were used as search models. Amino acid residues 320 to 406 were built and refined but electron density could not be observed for the first 2 and last residues most likely due to disorder. The structure was refined to 1.7Å enabling that all the DNA nucleotides and 111 water molecules could be modeled. The final, refined model (**Figure 3**) has an R_work_ of 20.7%, R_free_ of 24.1%, a B-factor of 49.8 and all backbone φ and ψ angles are within allowed regions. A full summary of the data collection and refinement statistics is shown in Table 1.

**Table 1:**
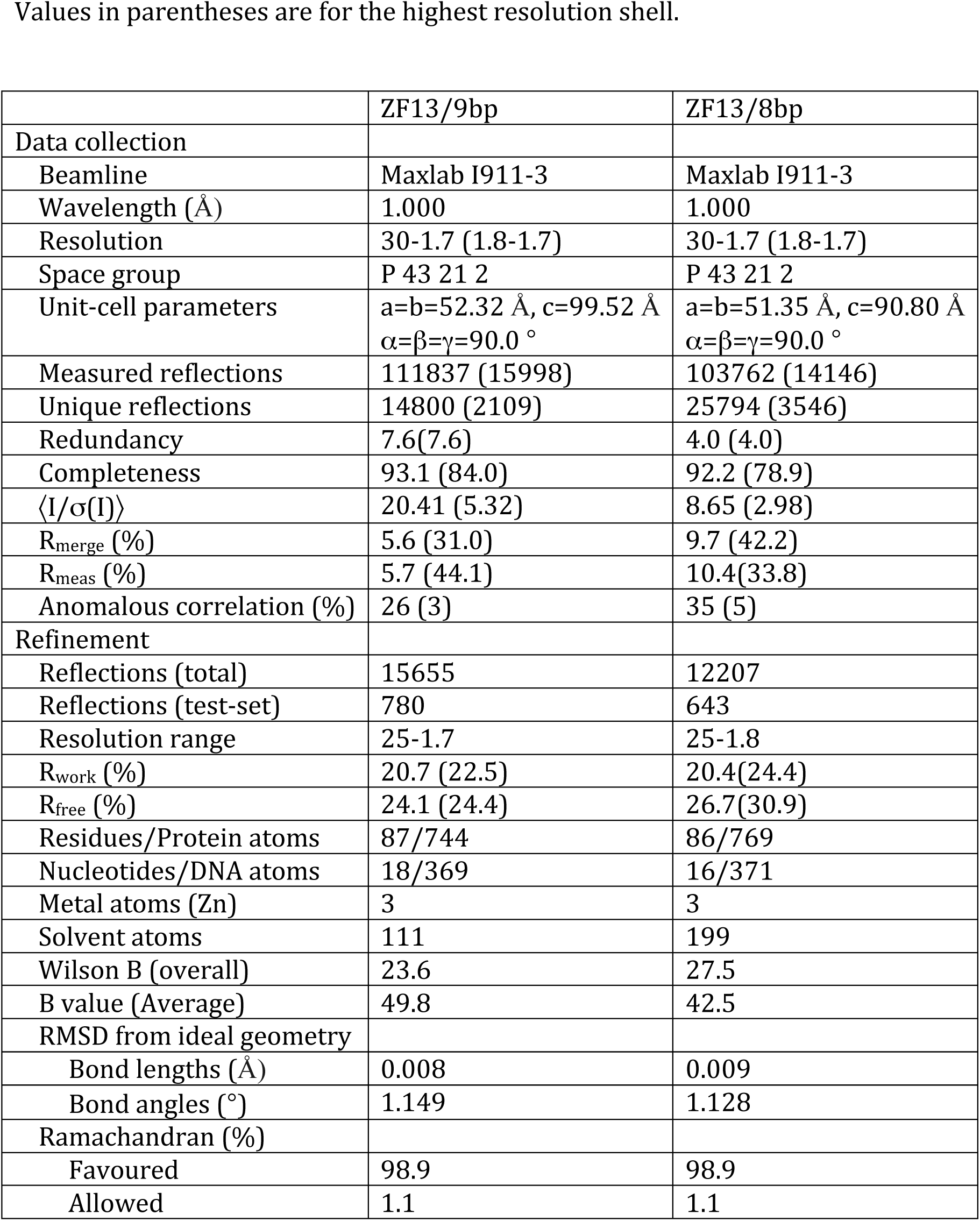
Data collection and refinement statistics Values in parentheses are for the highest resolution shell.

**Figure 3:**
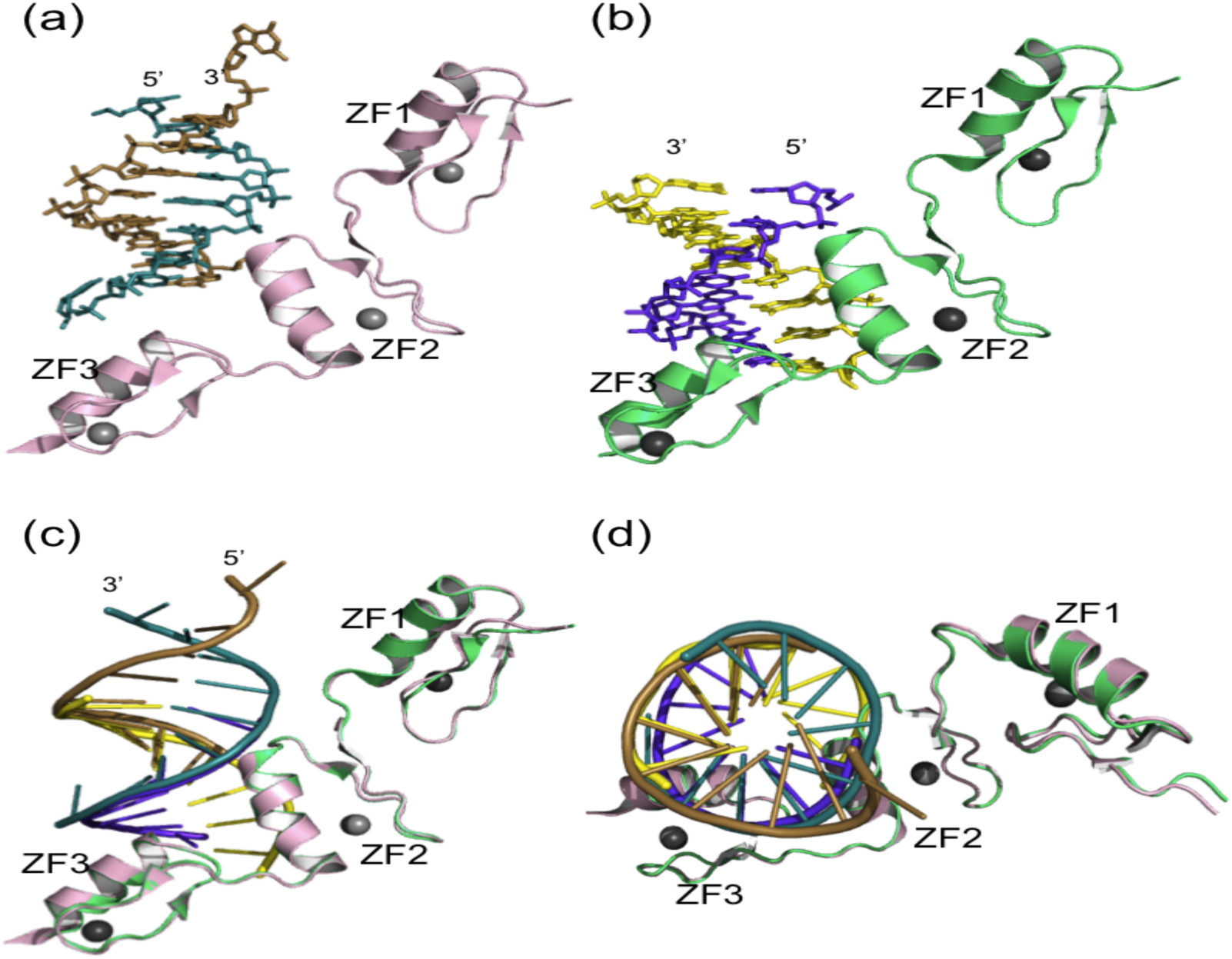
The individual refined structures and their superposition. (a) Structure of WT1 zinc finger 1 to 3 bound to a 9 bp DNA (ZF13/9bp). The coding stand is in sand, thenon-codingstrandinblue-greenandtheproteininlightpink.(b)Structure of WT1 zinc fingers 1 to 3 bound to an 8 bp DNA (ZF13/8bp). The coding strand is in yellow,the non-codingstrandinpurpleandtheproteinindarkgreen.Thecolour coding in (a) and (b) maintained throughout this report. (c) The superposed ZF13/9bp and ZF13/8bp structures viewed from the side and (d) viewed from the top, down the DNA double helix.

The first obvious feature of the ZF13/9bp structure (**Figure 3**) is the translocation of the DNA, placing it out of register so that each zinc finger does not bind at the expected base positions. Instead of binding at the relative DNA binding position 4-12, which provides DNA binding sites for ZF13, it binds at positions 8-15 only providing possible DNA binding triplet base-pairs for zinc finger 1 and 2 (**Figure 1b** and **Figure 4**). As a consequence, the DNA was renumbered to reflect its position in the crystal structure and all later references will use the new numbering (**Figure 2**). The structure shows however that the typical zinc finger fold is conserved for all 3 zinc fingers consisting of a β-hairpin loop followed by an α-helix held together by a Zn2+ ion coordinated by two cysteines and two histidine residues. Zinc fingers 2 and 3 adopt the conformation typical of C2H2 zinc finger domains in complex with DNA with the α-helix lying along the major groove with its N-terminal end dipped into the groove. Zinc finger 2 is perfectly placed in a major groove making all the expected contacts. Zinc finger 3 on the other hand is only held in place by a single hydrogen bond interaction with the backbone phosphate of C52 given that it usual binding site, base-pairs 4-7 is absent. The overall binding mode where the helix lies in the major groove is still conserved since the symmetry related molecule provides a binding site.

**Figure 4:**
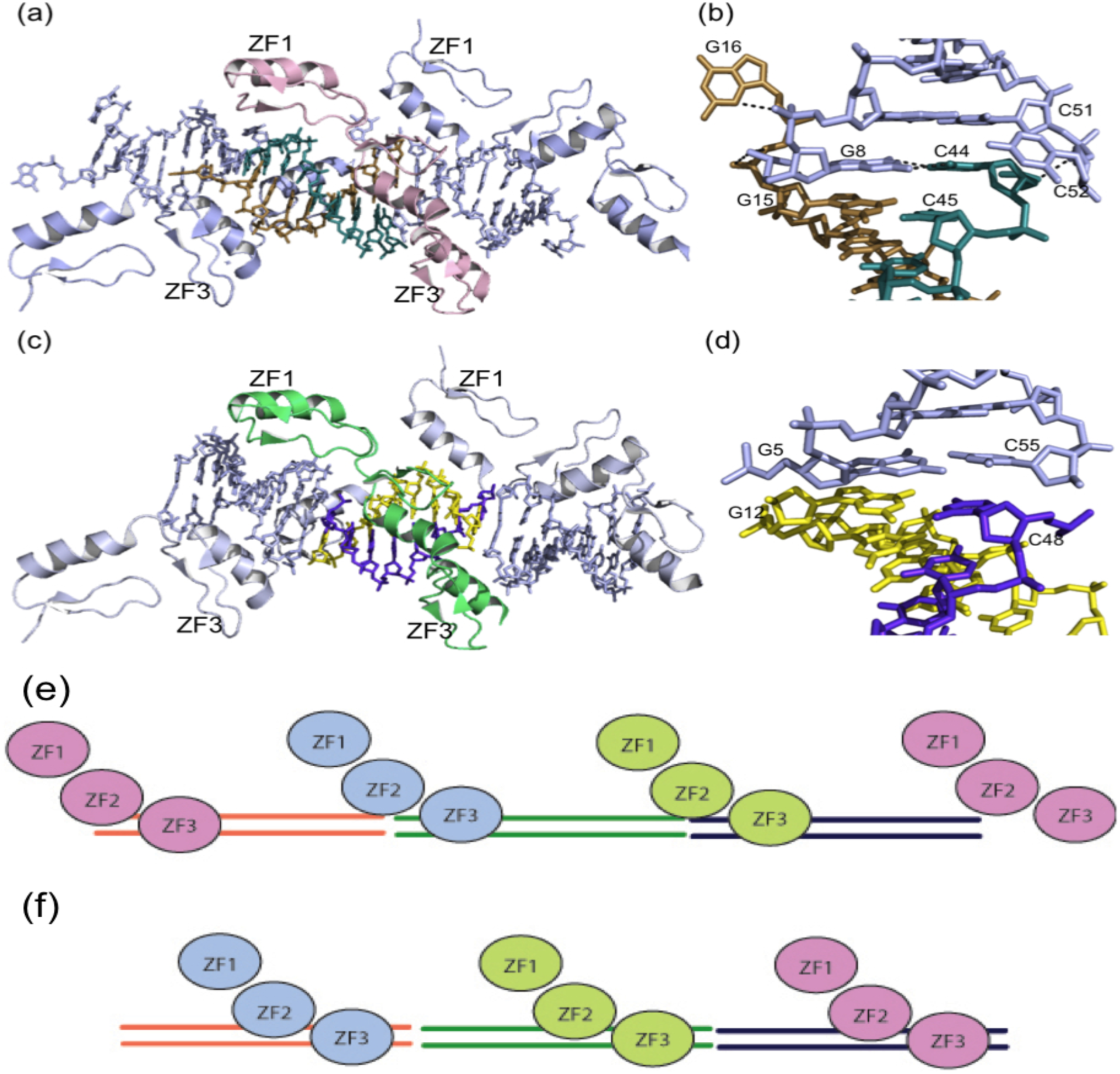
A representation of the arrangement of the zinc fingers and DNA in the structures. (a) and (c) show the arrangement of the complex molecules in the crystal of ZF13/9bp and ZF13/8bp respectively. Stacking one on the other to form a continuous DNA helix with tandem repeats of 3 zinc fingers wrapped around it. Adjacent molecules are coloured light blue and this is respected for the rest of this report. (b) and (d) show detailed packing of the DNA in the ZF13/9bp and ZF13/8bp structures respectively. (e) The binding position of the zinc fingers with respect to the DNA in the ZF13/9bp structure. (f) The expected binding positions of the zinc fingers to the DNA in both structures.

Generally the crystal packing interactions in this crystal are extensive with the individual molecules packed so closely that there is a very low solvent content of 41% in the crystal. Typical zinc finger DNA interactions and zinc finger 1 interactions mostly make the crystal packing interactions in this crystal lattice (**Figure 4**). However the DNA mediated crystal packing in this structure is unusual (**Figure 4**). The DNA packs head to tail with the 3’ nucleotide of each strand, G16 and C52 flipped out so that the 5’ nucleotides on opposite strands from the DNA in an adjacent molecules base-pair with each other. G8 base-pairs with C44 from the adjacent molecule instead of with the C52 from the non-coding strand in the same molecule. This effectively reduces the number of base-pairs per DNA helix in each molecule from 9 to 8. The base-pairing and stacking at this packing interface however respects the Watson-Crick type base-pairing mode. The base pairing and base stacking interactions at this interface are further stabilized by the stacking of the flipped out C52 base against the deoxyribose ring of C44 and 2 hydrogen bonds, one between the terminal phosphates of C44 and the backbone phosphate of the flipped out C52 and the other between the terminal phosphate of the flipped out G16 and the backbone phosphate of G8. There are other crystal packing interactions involving zinc finger 2 and 3. Zinc finger 2 packs against another zinc finger 2 from a neighboring molecule using the surface opposite to the DNA binding surface to make hydrophobic contacts using Try353, Gln374 and Thr378. This hydrophobic contact is flanked by hydrogen bonds, one between Lys371 and Thr378 on one side and one between Asp359 and Thr406 on the other side. Zinc finger 3 makes a single crystal contact, which is a hydrogen bond between Thr404 and Arg345 from zinc finger 2 in a neighboring molecule. The most significant crystal contacts are zinc finger DNA interactions mediated by zinc finger 2 and 3 where they recognize DNA from the adjacent molecule in the manner typical of C2H2 zinc fingers. The α-helix of each finger lies along the major groove of the DNA in the adjacent molecule and makes extensive base specific and backbone contacts. The linker region between zinc finger 2 and 3 and the interaction between the two fingers are identical to those in the previous structures of ZF24 bound to DNA [27, 33].

Zinc finger 1 in this structure does not bind at its expected DNA binding site. Instead, it is flipped out, away from its binding site making no contact with the DNA in this region. It rather extends out to pack against a symmetry related molecule inserted between the zinc finger 1 and DNA from the symmetry related molecule (**Figure 5**). The finger makes very strong hydrophobic interactions with zinc finger 1 from the adjacent molecule by means of a hydrophobic patch contributed by Phe324, Leu337, Leu340 and the side-chain ethyl of Glu341 all found on the N-terminal end of the α-helix and the first β-strand. This N-terminal end of the α-helix is the end that would normally interact with DNA. The hydrophobic patch from zinc finger 1 interacts with an identical hydrophobic patch from the adjacent zinc finger 1. On the opposite side of this hydrophobic interaction, zinc finger 1 interacts with DNA from the symmetry related molecule, mainly making backbone interactions including 2 hydrogen bonds, one between the side-chain of Tyr334 and the backbone phosphate of G9 and the other between the side chain of Lys332 and the same backbone phosphate of G9.

**Figure 5:**
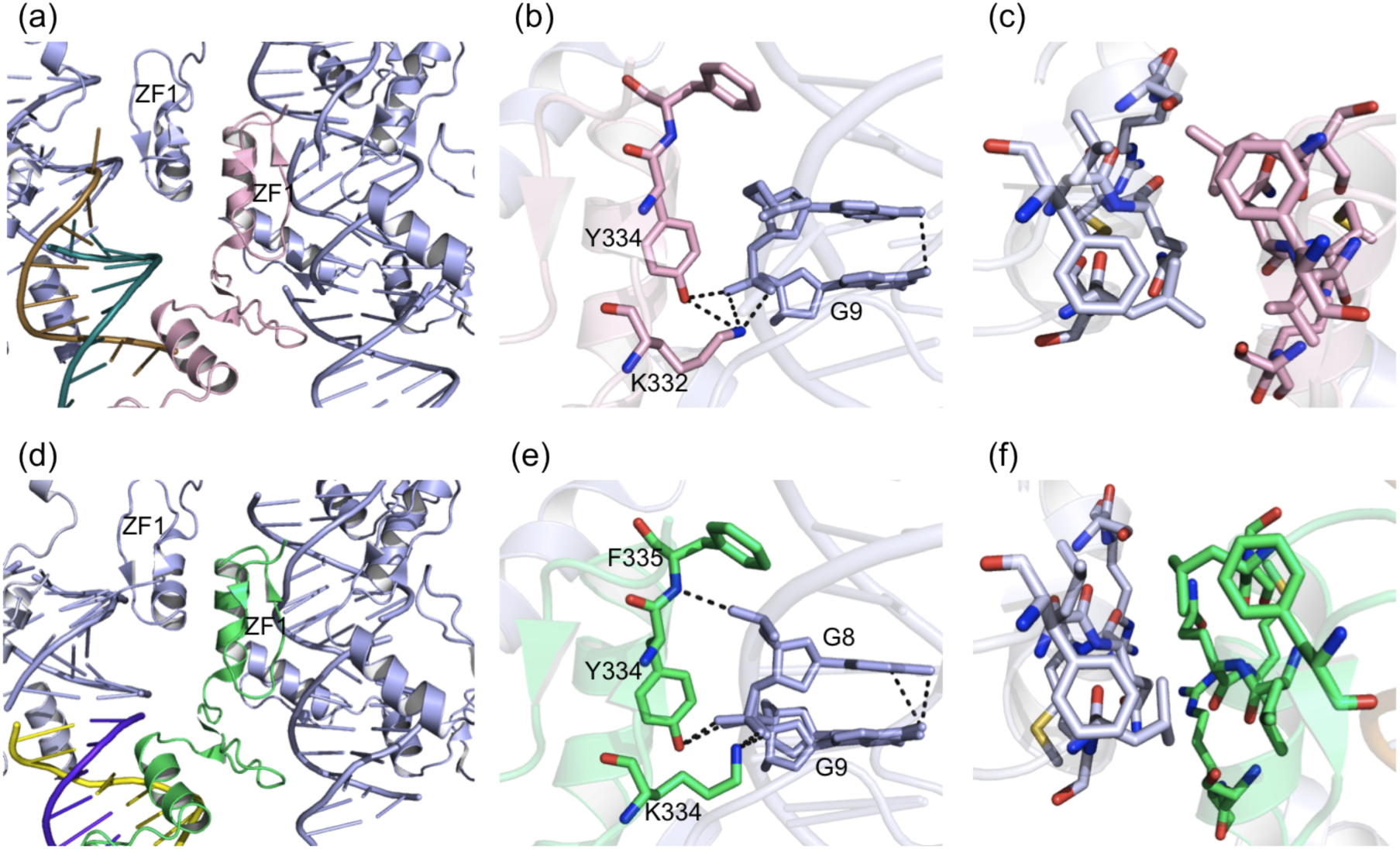
Binding mode of zinc finger 1 in the crystal structures. (a) and (b) show the interaction mode of zinc finger 1 in the ZF13/9bp and ZF13/8bp structures respectively. (b) and (d) show the precised hydrogen bond interactions made by zinc finger 1 with the DNA in the adjacent molecule in the ZF13/9bp and ZF13/8bp structures respectively. (c) and (f) show the hydrophobic interface between two zinc finger 1s from adjacent molecules in the ZF13/9bp and ZF13/8bp structures respectively.

### Structure of the WT1 ZF13 bound to its 8 bp cognate DNA (ZF13/8bp)

This complex was prepared as a result of the fact that in the ZF13/9bp structure, the zinc fingers seemed to be out of register. This, coupled with the fact that the 3’ nucleotide on each strand was flipped out resulting in an 8 bp long DNA helix and the fact that in previous structures of the ZF24-9bp and ZF24+9bp, the forth basepair, T4:A56 did not participate in the interaction rationalized the design of the ZF13/8bp complex. Data was collected on the crystals of this complex and indexed using the XDS program package in space group P4(3)2(1)2. The structure was solved by molecular replacement with 1 molecule per asymmetric unit and a solvent content of 40.8% using the ZF13/9bp structure as a search model. Amino acid residues 320 to 405 were built and refined but electron density could not be observed for the first 2 and last 2 residues. The structure was refined to 1.7Å enabling all the DNA nucleotides and 198 water molecules to be modeled. The final, refined model, presented in **Figure 3**, has an R_work_ of 20.4%, R_free_ of 26.7%, a B-factor of 42.5 and all backbone φ and ψ angels are within allowed regions. A full summary of the data collection and refinement statistics is also shown in table 1.

The global features of the ZF13/8bp structure are identical to those of the ZF13/9bp structure. Structure superposition reveals a global RMSD between all atoms in the two structure of only 0.47Å. The RMSD between all atoms in zinc finger 1 and in zinc fingers 2 and 3 is 0.42Å and 0.27Å respectively. This indicates that the protein in both structures is mostly identical with only very small variations, which are even smaller in zinc finger 2 and 3 compared to zinc finger 1. The main difference between the ZF13/9bp and the ZF13/8bp complexes is the relative position of the DNA. The DNA in the ZF13/8bp structure binds at the expected relative position providing the usual binding sites for zinc finger 1 and 2 and a partial binding site for zinc finger 3. Zinc finger 1 again does not bind DNA in this structure and it adopts the same position as in the ZF13/9bp structure. The DNA in the ZF13/8bp structure similarly packs in the crystal by stacking head to tail but no bases are flipped out since there are only 8 base-pairs in the fragment. The packing is such that the base stacking distance between the last base-pair in one molecule and the first in the adjacent molecule is at an ideal distance for Watson-Crick base stacking with G12 and C48 stacking against G5 and C55 respectively. Most of the other packing interactions mediated within this crystal are identical to those within the ZF13/9bp structure with a few exceptions. The 2 crystal packing hydrogen bonds mediated by zinc finger 2 in the ZF13/9bp structure are not formed in the ZF13/8bp structure due to an alternate conformation of Lys371 and the fact that Thr406 is not visible in the electron density. The one crystal packing hydrogen bond mediated by zinc finger 3 in the ZF13/9bp is also not formed in the ZF13/8bp structure also due to the alternate conformation of Arg345. The packing of zinc finger 1 in the ZF13/8bp structure is mostly identical to that in the ZF13/9bp structure but for an additional hydrogen bond formed between the backbone amide of Phe335 and the backbone phosphate of C8.

### Individual zinc finger interactions with DNA

Analyses of the zinc finger DNA binding interactions looking at only one molecule per asymmetric unit does not show most of the actual interactions mediated in the crystal forms obtained. This is due to the fact that the DNA from each molecule is effectively shared between zinc fingers from that molecule itself, as well as from an adjacent molecule. Analysis of one molecule is synonymous to analysis of a single hand in a firm handshake shared by two individuals, which cannot depict the whole picture. The DNA contacts mediated by the zinc fingers in the one molecule per asymmetric unit ZF13/9bp and the ZF13/8bp structures as calculated by Nucplot [42] (shown in supplementary material) show that zinc finger 3 makes only 1 backbone contact with the DNA which is not the actual case. To get a full picture of the DNA contacts made by each of the zinc fingers in the two structures, an analysis of the symmetry related molecules was also made. This analysis, which included the molecule in the asymmetric unit and the two symmetry related molecules making DNA stacking crystal packing interactions yielded a more complete picture of the zinc finger DNA contacts in the two structures. For a complete DNA interaction pattern for zinc finger 1 to 3, it was also necessary to do a comparative analysis of the two structures in other to incorporate some features of one structure to another since the DNA in each structure occupied a different register. This incorporation compensated for the differences in register and a difference in sequence as a result of that difference in register. The final results are models (**Figure 6**) that depict the complete DNA binding contacts observed in both the ZF13/9bp and ZF13/8bp structures. These two models diverge in the sense that the DNA represented in each structure differ from that in the other by 2 base-pairs which are simply flipped from either GC to CG or vice versa (**Figure 6 and 7**).

**Figure 6:**
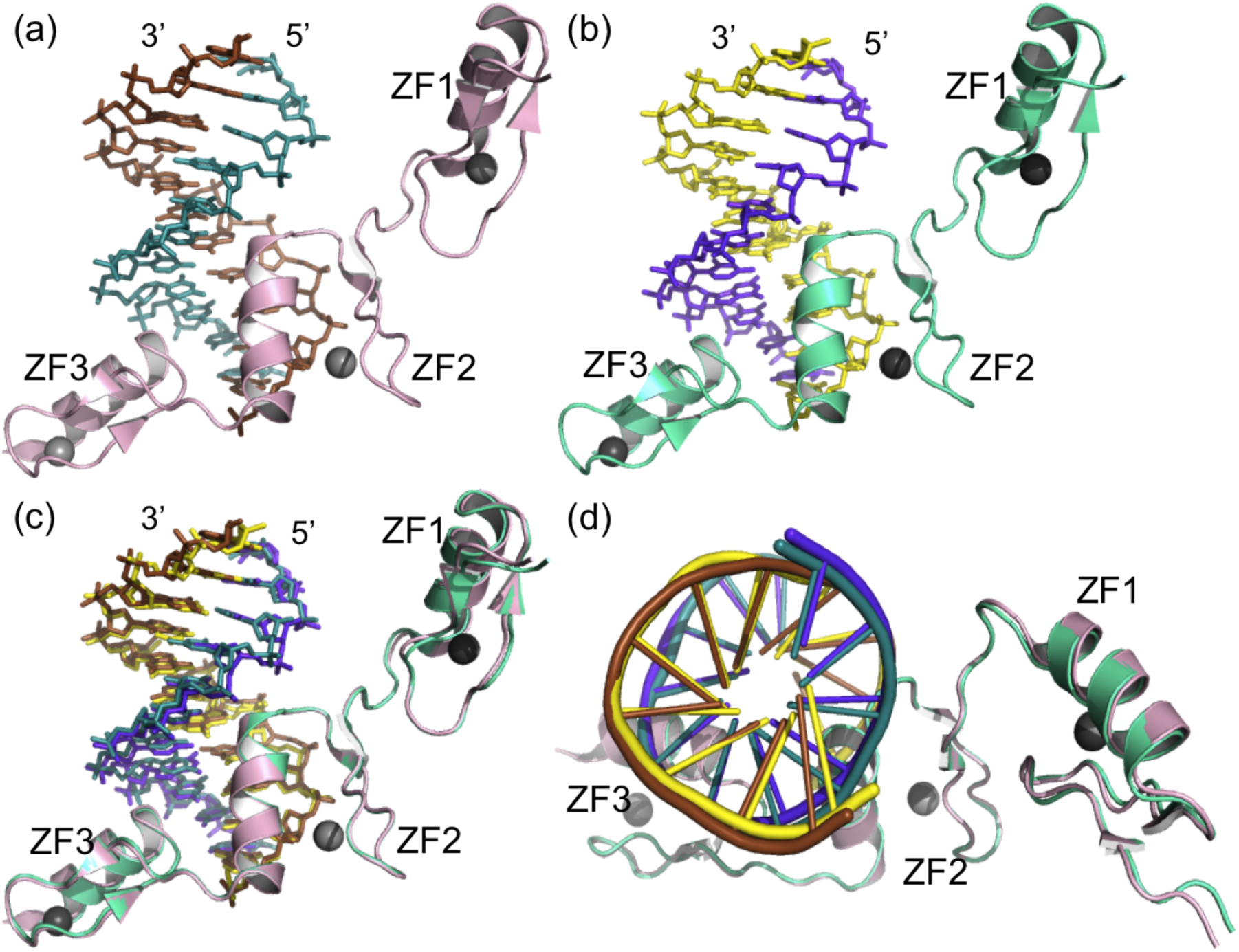
The same representation of the ZF13/9bp and ZF13/8bp structures as in figure 3 but the DNA in these structures has been remodeled based on analysis of the symmetry related molecules in the crystal and comparism of the two structures. The DNA in the structures of ZF13/9bp (a) and ZF13/8bp (b) is each 10 bp long. (c) and (d) show the true similarity in between the two structures when superposed, vied from the side and from the top respectively. These are the models used for the final DNA binding analysis.

**Figure 7:**
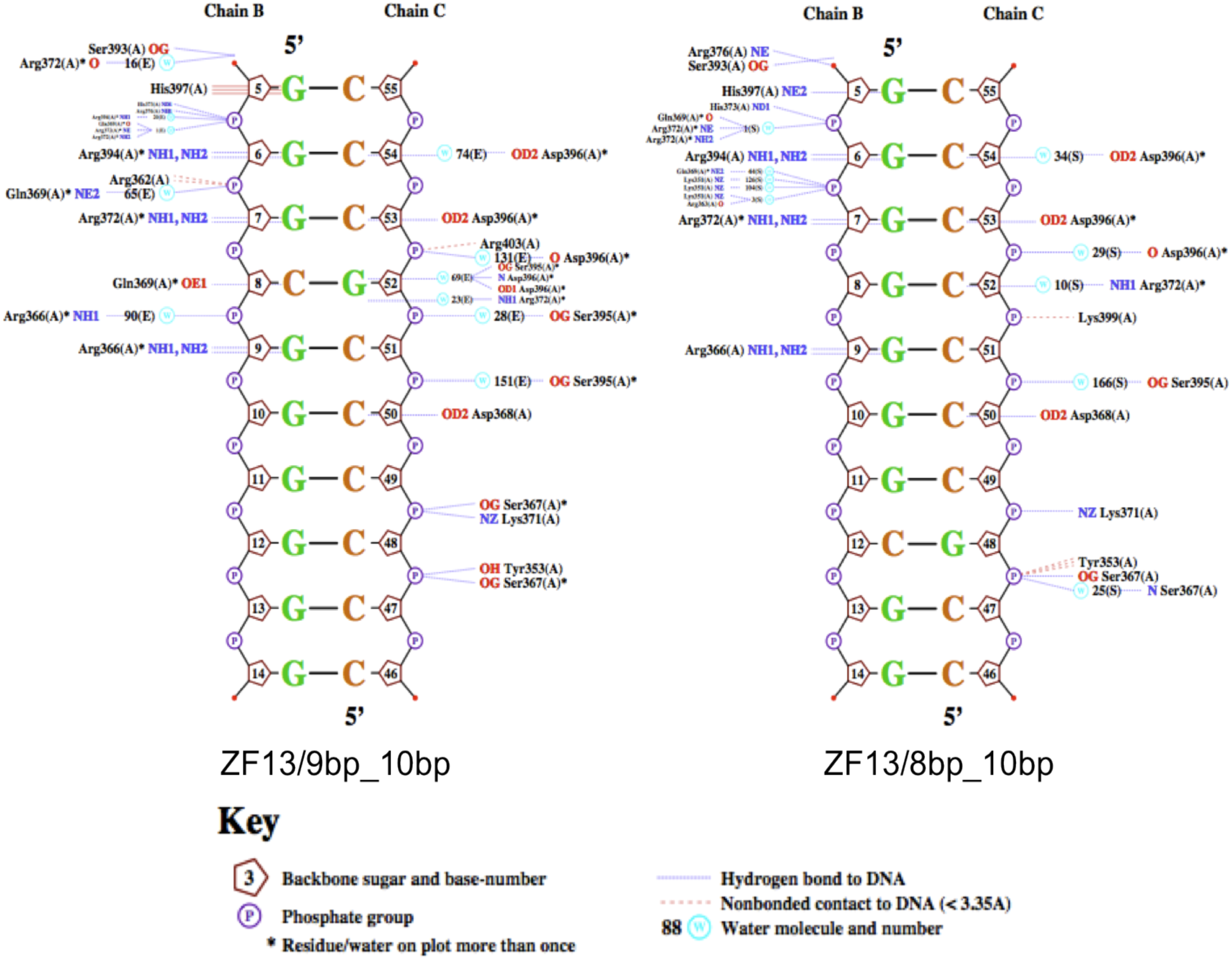
An overview of the interactions in the two structures as calculated by Nucplot.

C2H2 zinc fingers recognize and bind DNA using amino acids at specific positions with respect to the α-helix with the first amino acid at the N-terminal end of the helix referred to as 1, the amino acid preceding that as -1 and the amino acid following that as 2. These classical C2H2 zinc fingers normally use amino acids at positions -1, 2,3 and 6 to bind DNA by making base specific contacts with the DNA via the side chains of these amino acids [43]. Since zinc finger1 does not bind to its designed target DNA site in these two structures, only the base specific contacts made by zinc fingers 2 and 3 are described (**Figure 8**). Apart from interactions at the base pair at nucleotide position 8 with respect to the coding strand, where the nucleotides in the two strands are switched in the two structures, all other base specific contacts are maintained except for some water mediated contacts. There are however differences in the DNA backbone mediated, non-specific contacts mainly due to the differences in the water coordination or slight changes in side-chain conformations. The base specific contacts made by zinc finger 3 are identical. These interactions include 2 hydrogen bonds between Arg394 and G6, a hydrogen bond between Asp396 and C53 and a water-mediated hydrogen bond between Asp396 and C54. One of the variations is the hydrogen bond between His397 and G5 in the ZF13/9bp structure, which is replaced by non-bonded contacts in the ZF13/8bp structure. The other variation is a base specific, water mediated hydrogen bond contact in the ZF13/8bp structure, which is missing in the ZF13/9bp structure. The main differences are found in zinc finger 2 because the switched base-pair at position 8 is in the triplet base-pair DNA binding site for finger 2. The identical base specific contacts made by finger 2 include two hydrogen bonds between Arg366 and G9, two hydrogen bonds between Arg372 and G7, a hydrogen bond between Asp368 and C50 and a water-mediated hydrogen bond between Arg372 and C52. The only difference in the base specific contacts is a hydrogen bond formed between Gln369 and C8 in the ZF13/8bp structure which is missing in the ZF13/9bp structure due to the fact that there is a G at position 8 instead.

**Figure 8:**
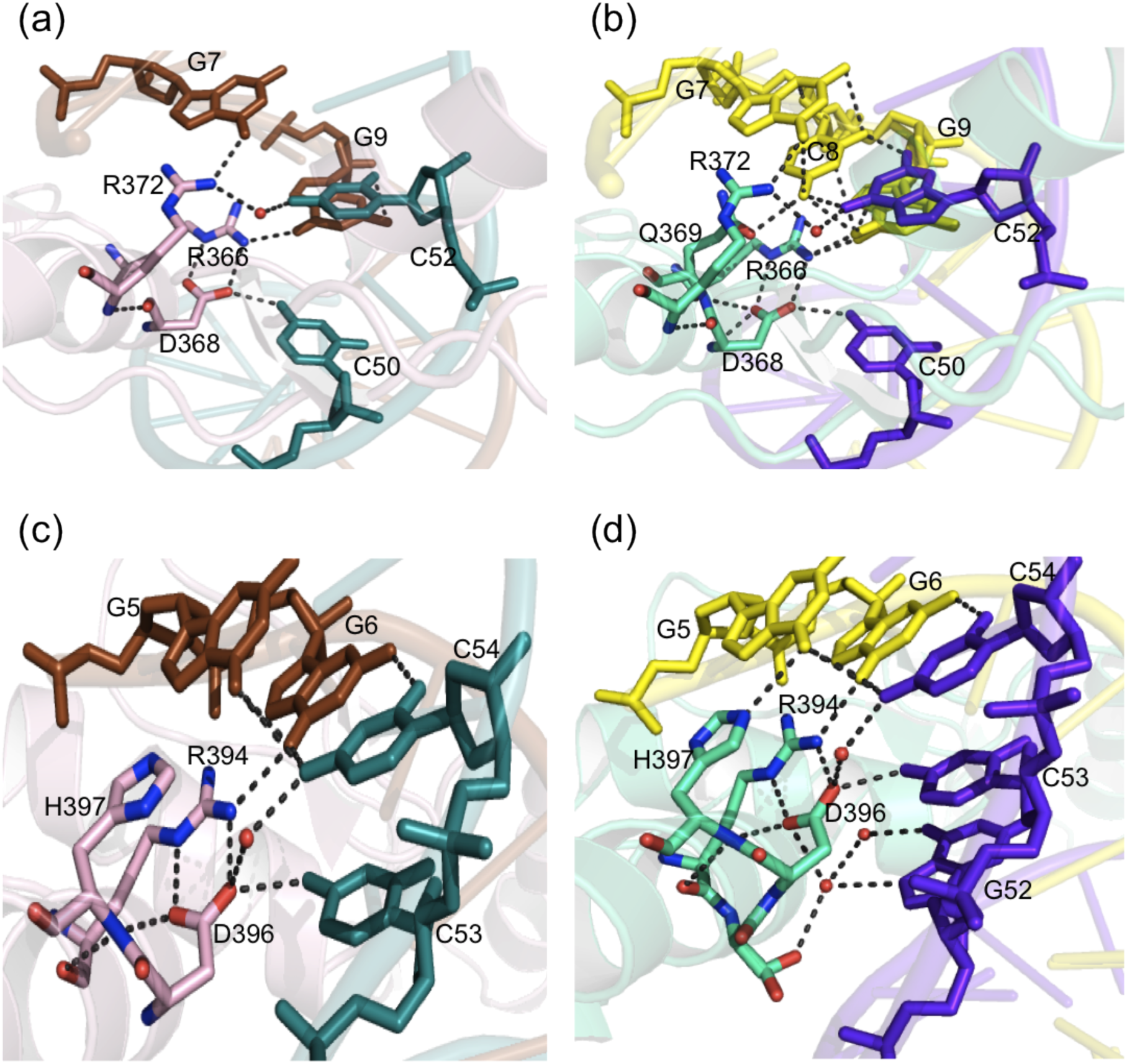
Comparison of the base specific interactions mediated by zinc finger 2 in the ZF13/9bp (a) and ZF13/8bp (b) structures. The same comparison is done for base specific interactions involving zinc finger 3, between the ZF13/9bp (c) and ZF13/8bp (d) structures.

## Discussion

Both the ZF13/9bp and the ZF13/8bp structures are refined to a high resolution of 1.7Å, which allows for a clear assignment of all the hydrogen bond interactions described. The average B factor of 49.8 and 42.5 for the ZF13/9bp and ZF13/8bp are at the higher average limit. This is however misleading, as the ribbon representation colored by B factor presented in **Figure 9** show that the average B factor for zinc finger 2 and 3 are quite low and only the average B factor for finger 1 and the frail ends of the DNA are high. This is consistent with the DNA binding results showing that zinc fingers 2 and 3 are bound to the DNA and fixed, therefore the low B factors. Zinc finger 1 which does not bind DNA and therefore is more flexible has high B factors. This is further observed in the electron density map, which is not as defined for zinc finger 1 as for zinc finger 2 and 3 in both structures. The apparent tendency for zinc finger 1 to be flexible in these highly packed structures may be an indication of *in vivo* flexibility intrinsic to zinc finger 1 in WT1 DNA interactions.

**Figure 9:**
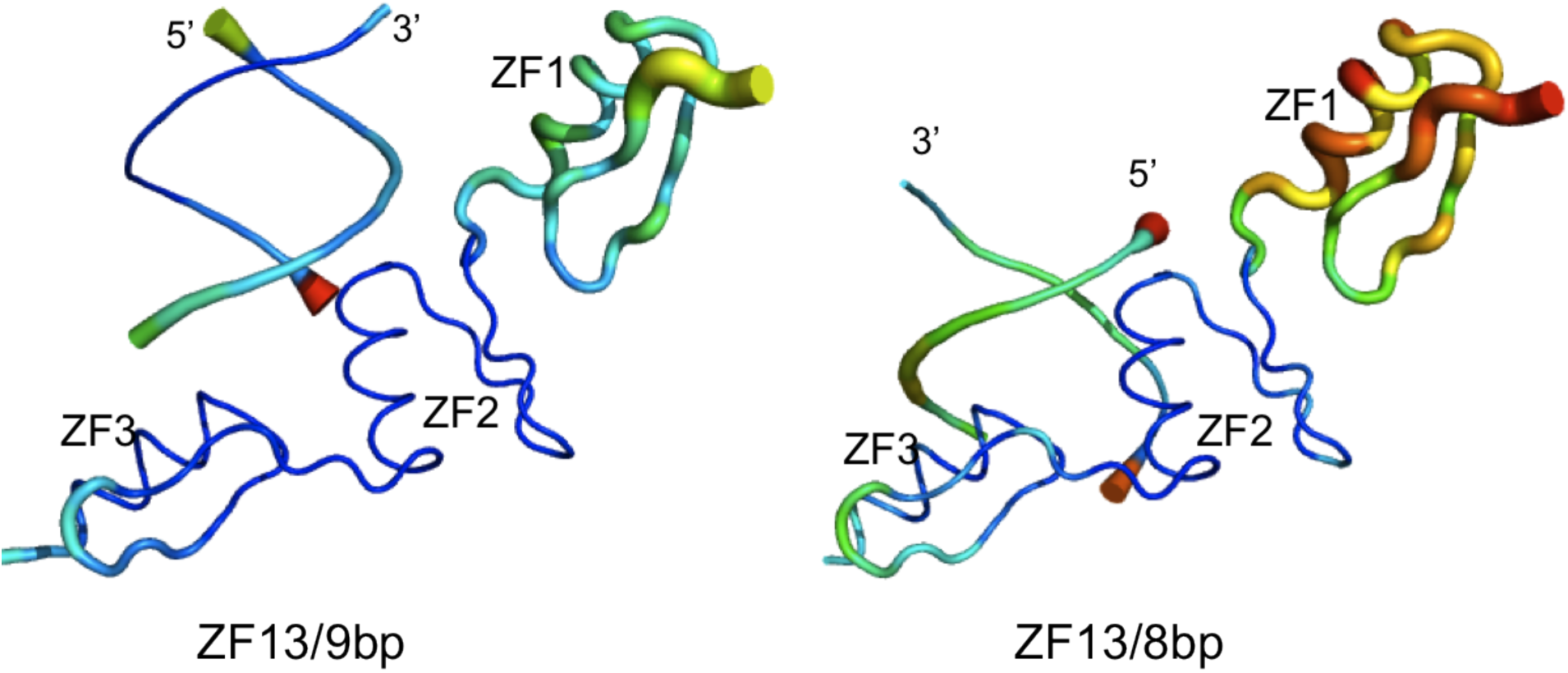
Ribbon representation of the ZF13/9bp and ZF13/8bp structures coloured according to the B factor. The larger the dimensions of the ribbon, the larger the B factor. The ribbon is colored from very low B factors in blue through medium B factors in light blue to high B factors in red.

The structures of zinc finger 2 and 3 bound to DNA in the ZF13/9bp and ZF13/8bp structures are consistent with the published crystal and NMR structures [27]. The present structures however reveal additional interactions that could not be observed in the previous structures due to limitations in resolution. The hydrogen bond assignments are more accurate and depict the actual hydrogen bond network in the WT1 zinc finger DNA interactions as opposed to the hydrogen bond network in the previous structures which was inferred from the structure of Zif268 zinc fingers bound to DNA [28]. There are some differences in the base specific contacts seen in the structures of ZF13/9bp and ZF13/8bp compared to the previous structures but not too many to disregard the previous findings. The additional non-specific backbone interactions and the water-mediated interactions, which have never been described before are complementary to the previous structures. Some of these water-mediated contacts are conserved between the ZF13/9bp and the ZF13/8bp structure and even mediate base specific contacts. Such new water mediated base specific contacts could have serious implications on the DNA specificity of the WT1 zinc finger domain, which could translate into a new understanding of its transcriptional activity. These structure show that ZF23 mediate very strong and highly specific interactions with DNA and therefore could act as the minimal DNA binding domain of WT1 as earlier proposed in a dual bacterial one hybrid and surface Plasmon resonance study [25].

The relative positions of the DNA in these two structures reveal a very fundamental aspect in zinc finger DNA interactions. Selectivity is more stringent in some positions, while other positions do not demonstrate any selectivity at all. As such, it is most often that transcriptional control elements have some flexible positions where more than one nucleotide can be tolerated. This has to be taken into consideration when searching gene promoter sequences to determine if they are possible transcriptional targets of a particular zinc finger transcription factor. It is known from previous selection studies that certain positions in the WT1 DNA binding sequence can tolerate substitutions at certain positions as demonstrated by the consensus sequences [22, 24, 26, 44]. The structural data presented here provide a molecular mechanism behind this selectivity. The selectivity for C at position 8 of the WT1 target DNA in the ZF13/8bp is as a result of a base specific hydrogen bond contact that is lost when this base is switched to a G in the ZF13/9bp structure. Such a loss of selectivity may not result in the complete lack of binding by the zinc finger domain but could still have serious consequences as has been demonstrated by the loss of a single base specific contact in the WT1 DNA interaction as a result of an Arg394Trp point mutation resulting in Denys-Drash syndrome [45]. These WT1 zinc finger domain point mutations that are linked to disease do not only disrupt base specific contacts but could also result in the de-stabilisation of the zinc finger resulting in the complete loss of binding activity by that zinc finger [45-48]. Even though most of the above mentioned point mutations are found in zinc fingers 2 and 3, it is clear from the structures that the specificity of zinc finger 4 is important for WT1 to effect transcriptional control on the correct genes. It will seem that without zinc finger 4, the WT1 zinc finger domain will sometimes recognize and bind to the wrong place on the genome as can be seen in the shift observed in the ZF13/9bp structure. Zinc finger 4 is therefore not only important, but vital to the correct function of WT1.

The particular role played by zinc finger 1 in DNA binding has up till now been unclear. While some evidence indicate that zinc finger 1 bind DNA firmly, similar zinc finger 2 and 3 and that it is indispensable in DNA recognition [22, 49], some suggest that it contributes very little to DNA binding and possibly acts as a protein or RNA recognition motif [24, 25, 30]. There has been some conflicting evidence on the exact way by which WT1 zinc finger one recognizes DNA. A study by Stoll and colleagues showed an X-Ray structure in which zinc finger one was partially bound to a distorted DNA strand, an event that would rarely happen in nature suggesting a crystallographic artifact. In the same study however, an NMR study structure show zinc finger one bound to DNA in a similar manner to the other zinc fingers [27]. In the ZF13/9bp and ZF13/8bp structures presented here, zinc finger 1 does not bind to its target DNA. It however makes some nonspecific contacts with the backbone phosphate of DNA from an adjacent molecule. These structures together show that zinc finger 1 does not have any specificity for DNA. Its interactions with the DNA backbone in all 4 structures indicate that it has some affinity for DNA. The fact that it adopts different positions in the two previously described structures compared to the present structures and that in the present structure it shows signs of flexibility despite some binding to the neighboring DNA suggest that its affinity for DNA is minimal and its binding to DNA is only transient and subject to proximity provided by the other zinc fingers that do bind DNA with high affinity and specificity.

Since evidence points to the fact that zinc finger 1 is not a dedicated DNA binding zinc finger, what possible role could it have? An RNA binding role has been described for the WT1 zinc finger domain with a central role for zinc finger 1 [30]. Other results have however shown that the other zinc fingers of WT1 still play a much more significant role in RNA interaction than zinc finger 1 [44, 50]. These conflicting evidences coupled with the fact that no real biological role has been described for WT1 RNA binding, apart from its co-localization with splicing factors in nuclear speckles, makes it premature to conclude that the WT1 zinc finger 1 is a biologically relevant RNA binding zinc finger. More evidence is needed to support the labeling of this zinc finger as an RNA binding zinc finger. In the present structure of ZF13/9bp and ZF13/8bp, zinc finger 1 also packs against another zinc finger by means of a hydrophobic patch located at the N-terminal end of the α-helix. This hydrophobic patch could be the reason for the lack of DNA binding by this zinc finger given that zinc finger DNA interactions are mostly based on electrostatic interactions depending on shape and surface charge complementarity. The interaction with the other zinc finger might be of functional relevance with WT1 zinc finger 1 having a functional role in protein-protein recognition. Some other DNA binding domains with multiple adjacent zinc finger topology such as the GLI, YY1 and TF111A have been shown to use 3 tandem zinc fingers for DNA recognition and others for protein-protein interactions [51-53].

The WT1 zinc finger domain recognizes DNA specifically by means of its zinc finger 2 to 4. Zinc finger 2 and 3 are responsible for most of the affinity and capable of acting as WT1 minimal DNA binding domain. Zinc finger 4 is however indispensible as it increases target specificity pinpointing a particular position on a target sequence for binding, preventing register promiscuity. While the biological role of the WT1 zinc finger 1 is still unclear, there is sufficient evidence to conclude that this zinc finger does not bind DNA specifically. It mediates a transient binding to DNA with minimal contribution to the overall DNA binding affinity of the entire domain. It may act as an RNA binding zinc finger or a protein binding zinc finger by means of hydrophobic interactions mediated by a hydrophobic patch on the α-helix.

